# Bidirectional changes of hepatic and peripheral insulin clearance characterize abnormal temporal patterns of serum insulin concentration in diabetic subjects

**DOI:** 10.1101/121392

**Authors:** Kaoru Ohashi, Masashi Fujii, Shinsuke Uda, Hiroyuki Kubota, Hisako Komada, Kazuhiko Sakaguchi, Wataru Ogawa, Shinya Kuroda

## Abstract

Insulin plays a central role in glucose homeostasis, and impairment of insulin action causes glucose intolerance and leads to type 2 diabetes mellitus (T2DM). A decrease in the transient peak and sustained increase of circulating insulin by glucose infusion accompany T2DM pathogenesis. However, the mechanism underlying this abnormal temporal pattern of circulating insulin concentration remains unknown. Here we show that bidirectional changes of hepatic and peripheral insulin clearance characterize this abnormal temporal pattern of circulating insulin concentration during the progression of T2DM. We developed a mathematical model using a hyperglycemic and hyperinsulinemic-euglycemic clamp in 111 subjects, including healthy normoglycemic and diabetic subjects. The hepatic and peripheral insulin clearance significantly increase and decrease, respectively, during the progression of glucose intolerance. The increased hepatic insulin clearance reduces the amplitude of circulating insulin concentration, whereas the decreased peripheral insulin clearance changes the temporal patterns of circulating insulin concentration from transient to sustained. These results provide insight that may be useful in treating T2DM associated with aberrant insulin clearance.

## Introduction

Insulin is the major anabolic hormone regulating the glucose homeostasis. The impaired action of insulin is a characteristics of type 2 diabetes mellitus (T2DM) (DeFronzo et al., 1992), accompanied by abnormality in the temporal patterns of circulating insulin concentration (Cerasi and Luft, 1967; Del Prato and Tiengo, 2001; Seino et al., 2011). In physiological condition, the circulating insulin concentration changes over the course of 24 h, including a persistently low level during fasting and a surge in response to food ingestion, which are known to be basal and additional secretions from the pancreas, respectively (Lindsay et al., 2003; Polonsky et al., 1988b).

Ability of additional insulin secretion is assessed by the oral glucose tolerance test (OGTT) (Stumvoll et al., 2000), in which a subject’s ability to tolerate the glucose load (glucose tolerance) is evaluated by measuring the circulating glucose concentration after an overnight fast (fasting plasma glucose concentration; FPG) and again 2 h after a 75-g oral glucose load (2-h post-load glucose concentration; 2-h PG) (Alberti and Zimmet, 1998). During this test, the circulating insulin concentration transiently increases and then continuously increases or decreases, known as the early and late phases of insulin secretion, respectively (Abdul-Ghani et al., 2006; Hayashi et al., 2013). The direct contribution of circulating glucose concentration to circulating insulin concentration is assessed by use of an intravenous glucose tolerance test (IVGTT) (Perley and Kipnis, 1967). This test excludes the effects of intestinal absorption of glucose and incretins secretion that trigger insulin secretion, thus permitting quantitative estimates of the ability of circulating glucose to initiate insulin secretion. During this test, the circulating insulin concentration transiently increases during the first 10 min and then continuously increases during the following 120 min, which are known as the first and second phases of insulin secretion, respectively (Pfeifer et al., 1981).

These temporal patterns of circulating insulin concentration differ across the stages of progression of T2DM. Based on an OGTT, a subject with FPG < 110 mg/dL (6.1 mM) and 2-h PG < 140 mg/dL (7.8 mM) is categorized as having healthy normal glucose tolerance (NGT). A subject with FPG of 110–125 mg/dL (6.1–6.9 mM) or 2-h PG of 140–199 mg/dL (7.8–11.0 mM) is categorized as having impaired glucose tolerance (IGT), and those with FPG ≥ 126 mg/dL (7.0 mM) or 2-h PG ≥ 200 (11.1 mM) as T2DM (Alberti and Zimmet, 1998). In general, plasma insulin concentration during the late-phase secretion of an OGTT in IGT subjects is higher than in NGT subjects, whereas the concentration during the early-phase secretion is similar in NGT and IGT subjects (Abdul-Ghani et al., 2006; Hayashi et al., 2013). Plasma insulin concentration during the first-phase secretion of an IVGTT decreases as glucose intolerance progresses, whereas that during the second-phase secretion is relatively maintained (Cerasi and Luft, 1967; Del Prato and Tiengo, 2001; Seino et al., 2011). Such changes of the temporal patterns of circulating insulin concentration during the progression of glucose intolerance from NGT to T2DM suggest these temporal patterns are involved in the maintenance and impairment of glucose homeostasis. Together with the measurement of circulating glucose concentration, the time course of circulating insulin concentration are used to assess insulin secretion from the pancreas and insulin sensitivity.

However, it is difficult to assess the insulin secretion and sensitivity of body tissues directly from the circulating insulin concentration because of the negative feedback between circulating insulin and glucose. A rise in circulating glucose concentration stimulates insulin secretion, and the resultant rise in circulating insulin concentration stimulates glucose uptake, causing circulating glucose concentration to fall. This feedback means there is mutual dependence between glucose and insulin, making it difficult to distinguish the effect of insulin secretion and sensitivity directly from the circulating insulin concentration (DeFronzo et al., 1979).

To directly assess insulin secretion without the effect of the feedback from insulin to glucose, DeFronzo et al. (1979) developed the hyperglycemic clamp technique, in which insulin secretion is measured while circulating glucose concentration is at a fixed hyperglycemic plateau maintained by exogenous continuous glucose infusion. The measurements of circulating insulin concentration during the first 10 min and after 10 min are used to assess the insulin secretion ability and are known as the first and second phase insulin secretions, respectively (DeFronzo et al., 1979; Okuno et al., 2013).

Conversely, to directly assess insulin sensitivity without the effect of the feedback from glucose to insulin, the hyperinsulinemic-euglycemic clamp was developed (DeFronzo et al., 1979). In this method, circulating insulin concentration is maintained at a fixed hyperinsulinemic plateau and circulating glucose at a fixed normal plateau by continuous infusion of both insulin and glucose. Tissue insulin sensitivity is defined as the ratio of the glucose infusion rate to the circulating insulin concentration when they reach plateaus (DeFronzo et al., 1979; Okuno et al., 2013).

The body controls the circulating insulin concentration by balancing insulin secretion and insulin clearance. The major organs responsible for insulin clearance are the liver, which removes portal insulin during first-pass transit (Duckworth et al., 1988; Sato et al., 1991), and insulin-sensitive tissues such as muscle, which remove insulin from the systemic circulation (Rabkin and Kitaji, 1983). The insulin clearance in the liver and in other organs is called hepatic and peripheral insulin clearance, respectively. Although the relationship between changes of insulin clearance and the progression of glucose intolerance have been reported, the effects of insulin clearance are controversial. Some studies found that during the progression of glucose intolerance, insulin clearance decreased (Jones et al., 1997; Lee et al., 2013; Marini et al., 2014; Rudovich et al., 2004), whereas hepatic insulin clearance increased (Tamaki et al., 2013) or decreased (Bonora et al., 1983; Marini et al., 2014). Thus, the hepatic and peripheral insulin clearances were not explicitly distinguished, making it difficult to interpret the effect of both types.

Hepatic insulin clearance cannot be assessed directly from circulating insulin concentration because insulin is extracted from the liver before secreted insulin is delivered into the systemic circulation. However, insulin is secreted at an equimolar ratio with C-peptide, a peptide cleaved from proinsulin to produce insulin, which is not extracted in the liver. Thus, by measuring circulating C-peptide concentration simultaneously with circulating insulin concentration, the pre-hepatic insulin concentration can be accurately assessed. The C-peptide index, which is the ratio of circulating glucose to C-peptide concentration, is an index of insulin secretion with clinical utility (Saisho et al., 2011). Hepatic insulin clearance is clinically quantified as the ratio of circulating insulin to C-peptide concentration during the first 10 min under the hyperglycemic clamp condition (Ahrén and Thorsson, 2003).

The clinical indices of insulin secretion and clearance are indirect measures because they are obtained from temporal patterns of circulating concentrations, which are simultaneously affected by insulin secretion and clearance. Therefore, the clinical index of insulin secretion implicitly involves the effect of insulin clearance and vice versa. Mathematical models have been developed for specifically quantifying insulin secretion, sensitivity, and clearance abilities from temporal patterns of circulating concentration by accounting for this mutual dependence (Breda et al., 2001; Cobelli et al., 2014; Picchini et al., 2005). The model known as the minimal model is used to estimate insulin sensitivity and insulin secretion abilities for each individual based on the time courses of circulating glucose and insulin concentrations during IVGTT (Bergman et al., 1981). Furthermore, from the parameters of the model, Bergman et al. identified a relationship between the subject’s glucose intolerance and the product of insulin secretion and sensitivity.

We previously developed a mathematical model based on time courses of plasma glucose and serum insulin during consecutive hyperglycemic and hyperinsulinemic-euglycemic clamp conditions, and estimated the parameters of insulin secretion, sensitivity, and peripheral insulin clearance for each subject. We found that peripheral insulin clearance significantly decreased from NGT to IGT to T2DM during the progression of glucose intolerance (Ohashi et al., 2015). However, the hepatic and peripheral insulin clearance could not be distinguished because C-peptide was not incorporated in the model (Ohashi et al., 2015).

Hepatic insulin clearance is calculated as the difference between pre-hepatic and post-hepatic insulin concentrations assessed by comparing circulating C-peptide and insulin concentrations, because C-peptide, unlike insulin, is not removed by the liver. Since the circulating C-peptide concentration is also controlled by its secretion and clearance, a mathematical model for C-peptide kinetics was developed (Eaton et al., 1980). The models for circulating insulin and C-peptide have been used to estimate the secretion and kinetics of insulin and C-peptide, as well as hepatic insulin clearance (Chase et al., 2010; Cobelli and Pacini, 1988; Grodsky, 1972; Ličko and Silvers, 1975; Mari et al., 2002; Toffolo et al., 2001; Toffolo et al., 2006; Vølund et al., 1987). However, peripheral insulin clearance was not assessed in the models, because exogenous insulin infusion, which is required for accurate estimation of peripheral insulin clearance, was not performed.

Recently, Polidori et al. (2016) reported that both hepatic and extrahepatic insulin clearance, corresponding to peripheral insulin clearance, can be estimated by modeling analysis using plasma insulin and C-peptide concentrations obtained from the insulin-modified frequently sampled IVGTT. The parameters of hepatic and peripheral insulin clearance in the model were not highly correlated, suggesting that the two types of insulin clearance are regulated differently. In addition, hepatic insulin clearance was negatively correlated with insulin secretion, and peripheral insulin clearance was positively correlated with insulin sensitivity. However, hepatic and peripheral insulin clearance in T2DM subjects and the roles of both types of clearance in the changes in temporal pattern of circulating insulin concentration during the progression of glucose intolerance have yet to be examined.

In this study, we developed a mathematical model based on the time course of the serum insulin and C-peptide concentrations during consecutive hyperglycemic and hyperinsulinemic-euglycemic clamp conditions, and estimated hepatic and peripheral insulin clearance for each subject. The parameters from 111 subjects (47 NGT, 17 IGT, and 47 T2DM) showed a significant increase in hepatic insulin clearance and significant decrease in peripheral insulin clearance from NGT to IGT and T2DM, respectively. We also found that hepatic and peripheral insulin clearance play distinct roles in the abnormal temporal patterns of serum insulin concentration during the progression of glucose intolerance, namely an increase in hepatic insulin clearance reduces the amplitude of serum insulin concentration, whereas a decrease in peripheral insulin clearance changes the temporal patterns of serum insulin concentration from transient to sustained.

## Results

### Consecutive hyperglycemic and hyperinsulinemic-euglycemic clamp data

We calculated the averaged time courses of concentrations of plasma glucose and serum insulin and C-peptide during consecutive hyperglycemic and hyperinsulinemic-euglycemic clamp conditions of NGT (*n* = 50), IGT (*n* = 18), and T2DM (*n* = 53) (*Figure 1; Figure 1-figure supplement 1*) (Ohashi et al., 2015; Okuno et al., 2013). During the hyperglycemic clamp, plasma glucose concentrations at the hyperglycemic plateau were almost similar among the NGT, IGT, and T2DM groups.

**Figure 1.**
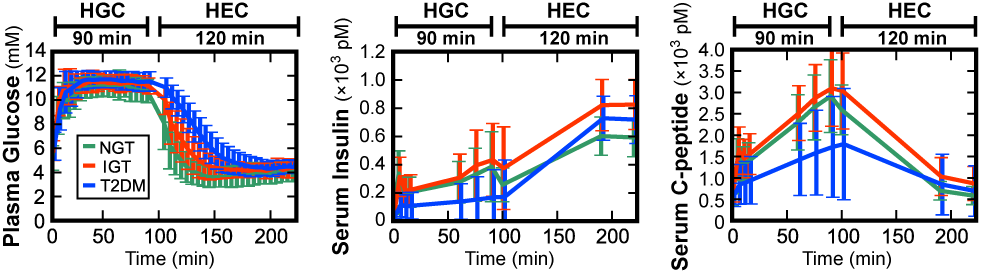
Concentrations of plasma glucose and serum insulin and C-peptide during consecutive hyperglycemic and hyperinsulinemic-euglycemic clamps. The mean ± SD among the subjects for NGT (green, *n* = 50), IGT (red, *n* = 18), and T2DM (blue, *n* = 53) are shown. Hyperglycemic clamp (HGC) was performed for 90 min and hyperinsulinemic-euglycemic clamp (HEC) for 120 min with a 10-min interval. The plasma glucose level is the average value calculated every 5 min of the measurements made every 1 min, and the serum insulin and C-peptide levels are measured values at sampling time (Materials and Methods). *Figure 1-source data 2* and *Figure 1-figure supplement 1* illustrate the significant difference of concentrations at each time point among the three groups.

Both the first (0–15 min) and second phase of insulin secretion (15–90 min) were clearly observed in the NGT and IGT subjects, whereas the two phases of insulin secretion were significantly reduced in the T2DM subjects. Serum C-peptide concentration showed a similar increase during the first and second phases of insulin secretion in the NGT and IGT subjects, whereas serum C-peptide concentration was significantly lower in the T2DM subjects during both phases. Although insulin and C-peptide should be secreted in an equimolar manner, the serum C-peptide concentration was higher than the serum insulin concentration because insulin—but not C-peptide—was removed by the liver and C-peptide clearance in the periphery was slower than insulin clearance.

During the hyperinsulinemic-euglycemic clamp at 100–220 min, serum insulin concentration was at a steady-state plateau of hyperinsulinemia, but serum insulin concentration differed significantly among the NGT, IGT, and T2DM subjects. The average serum insulin concentration of the NGT subjects was lowest and that of the IGT subjects was highest. These differences indicate that the ability to remove infused insulin from serum is different among the three groups and suggest that the difference lies in the peripheral insulin clearance. The plasma glucose concentration returned to the basal level from hyperglycemia at a different decay rate among the three groups. The average decay rate was lowest in the T2DM subjects and highest in the NGT subjects, suggesting that insulin sensitivity, which is the ability to promote the hypoglycemic effect in response to serum insulin, decreases during the progression of glucose intolerance. The serum C-peptide concentration returned to the fasting level in all groups. Only insulin was infused during the hyperinsulinemic-euglycemic clamp, indicating that serum C-peptide was derived only from endogenous secretion.

### Mathematical model for serum insulin and C-peptide concentrations

Many mathematical models that reproduce circulating insulin and C-peptide concentrations have been developed (Bergman et al., 1981; Cobelli and Pacini, 1988; Grodsky, 1972; Ličko and Silvers, 1975; Toffolo et al., 2001; Toffolo et al., 2006; Vølund et al., 1987). We developed six mathematical models based on these models, and the best model was selected for reproducing measured serum insulin and C-peptide concentrations during consecutive hyperglycemic and hyperinsulinemic-euglycemic clamp (Ohashi et al., 2015; Okuno et al., 2013) (*Figure 2-figure supplement 1*). These models have serum insulin and C-peptide concentrations including both insulin and C-peptide secretion and their hepatic and peripheral clearance. Plasma glucose perturbation and insulin infusion were used as inputs. For each of the 121 subjects, parameters of the six models were estimated by using measured concentrations of plasma glucose and serum insulin and C-peptide. We employed the model to reproduce serum insulin and C-peptide time courses with a simpler model structure, which took the minimum value of Akaike information criterion (AIC) in the six mathematical models (Akaike, 1973). The AIC is used as a criterion of the goodness of fit of a model.

The model consisting of four variables (*Model VI* in *Figure 2-figure supplement 1*) was selected as the best model with the minimum AIC for 76 of 121 subjects (*Figure 2A, Table 1*). In this model, the variables *I* and *CP* correspond to serum concentrations of insulin and C-peptide, respectively. The variable *X* corresponds to stored insulin and C-peptide in β-cells or β-cell masses. Because the amounts of stored insulin and C-peptide are equal, a single variable, *X*, is used for both. The variable *Y* is the insulin provision rate depending on plasma glucose concentration. The differential equations of the model are as follows:

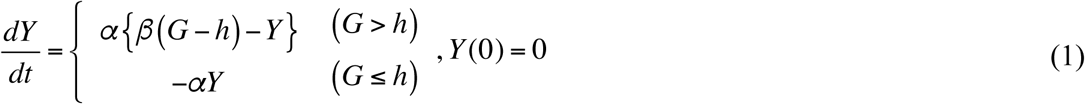

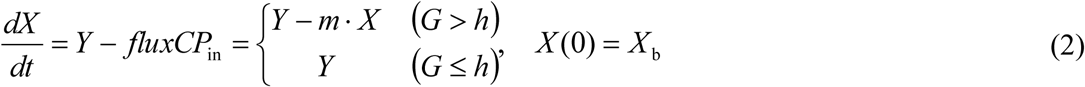

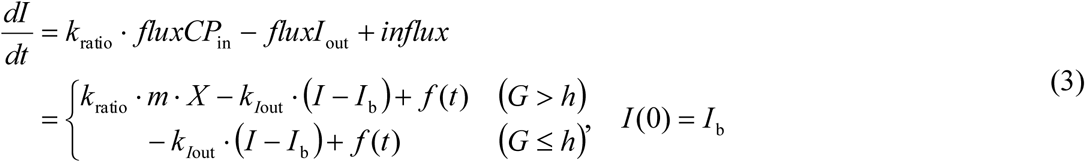

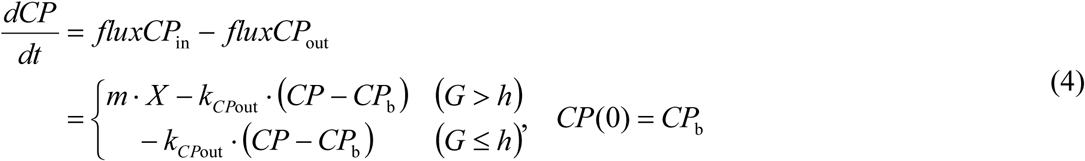
where *I_b_* and *CP_b_* correspond to fasting (basal) serum insulin and C-peptide concentration, respectively, directly given by the measurement, and *X_b_* is an initial value of *X* to be estimated.

**Figure 2.**
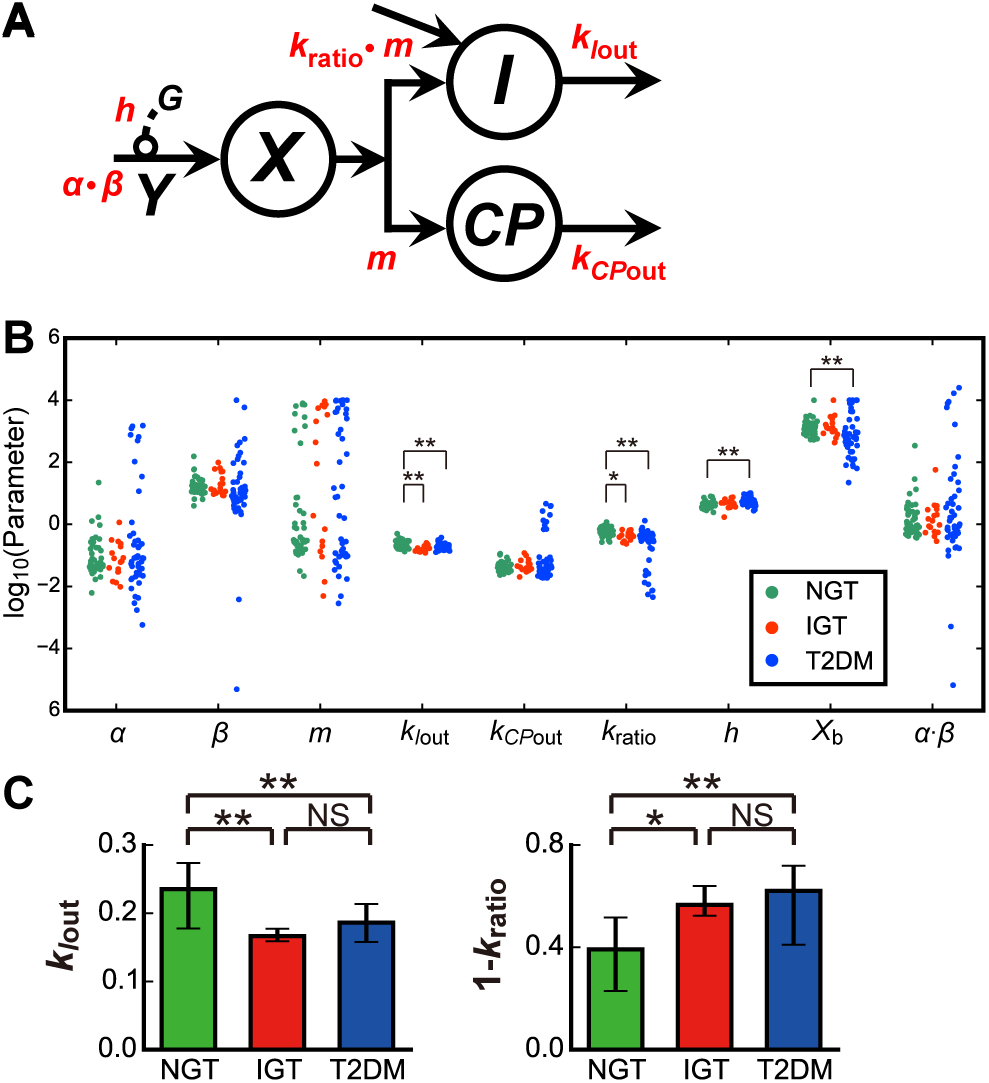
Mathematical model of serum insulin and C-peptide. (A) The structure of the model (see also Eqs. 1–4 and *Model VI* in *Figure 2-figure supplement* 1)*.I* and *CP* are serum insulin and C-peptide concentration, respectively. *X* is the amount of stored insulin and C-peptide, and *Y* is provision rate controlled by plasma glucose concentration, *G*. Arrows indicate fluxes with corresponding parameters (red). (B) The estimated parameters for the NGT (green), IGT (red), and T2DM (blue) subjects. Each dot corresponds to the indicated parameter for an individual subject. (C) The averaged parameters of *k*_*I*out_ and (1 – *k*_ratio_), corresponding to peripheral and hepatic insulin clearance, respectively. \**P* < 0.05, \*\**P* < 0.01, NS: not significant (Streel-Dwass test).

**Table 1.** Comparison of the models based on the Akaike information criterion (AIC)

| Model | No. subjects<br>of min AIC | NGT | IGT | T2DM | AIC<br>mean ± SD |
| --- | --- | --- | --- | --- | --- |
| *I* | 25 | 7 | 4 | 14 | $-21.1 \pm 22.9^*$ |
| *II* | 5 | 3 | 1 | 1 | $-19.0 \pm 24.2^*$ |
| *III* | 0 | 0 | 0 | 0 | $-16.2 \pm 24.9^*$ |
| *IV* | 7 | 0 | 0 | 7 | $-17.2 \pm 25.3^*$ |
| *V* | 8 | 4 | 3 | 1 | $-28.4 \pm 26.2$ |
| *VI* | 76 | 36 | 10 | 30 | $-31.4 \pm 25.2$ |
| Total | 121 | 50 | 18 | 53 |  |

The number of subjects optimal for each model with minimum AIC is shown (see Materials and Methods). \*AIC different from *Model VI* ( $P < 0.01$ , corrected by the number of *t*-tests, multiplied by 5).

Eq. 1 describes how insulin provision rate *Y* increases according to *αβ*(*G – h*) when *G* > *h*, otherwise zero, and decreases with *αY*. This means that provision of *X*, stored amounts of insulin and C-peptide, is linearly related via a time constant 1/*α* and parameter *β*, and stimulated only when the plasma glucose concentration exceeds the threshold value, *h*, which may correspond to the fasting plasma glucose concentration.

Eq. 2 describes how *X* increases according to the provision rate *Y* and decreases according to the insulin and C-peptide secretion *flux CP*_in_. *flux CP*_in_ is *X* secreted at the rate *m* when *G* > *h*, otherwise zero.

Eq. 3 describes how serum insulin concentration *I* increases according to the post-hepatic insulin delivery, *k*_ratio_ · *flux CP*_in_, and infused insulin, *influx*, and decreases according to peripheral insulin clearance *flux I*_out_. *k*_ratio_ · *flux CP*_in_ is expanded as *k*_ratio_ · *m* · *X*, which corresponds to insulin delivered into peripheral circulation after passage through the liver, when *G* > *h*, otherwise zero. The parameter *k*_ratio_ is the molar ratio of post-hepatic insulin to C-peptide, which represents the fraction of insulin delivered to the peripheral circulation without being extracted by the liver. Given that C-peptide is not extracted by the liver, *k*_ratio_ can represent the remaining fraction of insulin after the extraction by the liver over the total amount of secreted insulin, and ranges from 0 to 1. Therefore, (1 – *k*_ratio_) represents the fraction of insulin extracted by the liver and not delivered to the peripheral circulation and corresponds to hepatic insulin clearance. *influx* is the insulin infusion rate during hyperinsulinemic-euglycemic clamp. The infusion rate at time *t* is represented by the function *f*(*t*) (Materials and Methods). *flux I*_out_ represents serum insulin degradation with the rate parameter *k*_*I*out_. Therefore, *k*_*I*out_ represents insulin degradation in the periphery and corresponds to peripheral insulin clearance.

Eq. 4 describes how serum C-peptide concentration *CP* increases according to the C-peptide secretion *flux CP*_in_ and decreases according to peripheral C-peptide clearance *flux CP*_out_. *flux CP*_in_ is C-peptide secreted and delivered to peripheral serum without hepatic clearance. *flux CP*_out_ represents serum C-peptide degradation with the rate parameter *k*_*CP*out_.

For the 45 of 121 subjects who were not optimal for this model (*Figure 2A; Model VI* in *Figure 2-figure supplement 1*), the distributions of the residual sum of squares (RSS) between modeled and measured concentrations of insulin and C-peptide in this model were not significantly different from the distributions of the RSS of the rest of 76 subjects who were optimal with minimum AIC for this model (*Figure 2-figure supplement 2*). There also seems to be no bias in the distribution of the NGT, IGT, and T2DM subjects among the best models (*Table 1*). However, in the RSS distributions of all 121 subjects in this model, RSS values of three subjects were relatively high and detected as outliers, and the three subjects were excluded from the analysis in this study (*Figure 2-figure supplement 2*). In addition, there seems to be no bias of temporal patterns of serum insulin and C-peptide concentrations among the subjects in each model (*Figure 2-figure supplement 3; Figure 2-source data 2*). Therefore, we selected the model (*Figure 2A; Model VI* in *Figure 2-figure supplement 1*) for further study because it was able to reproduce time courses of serum insulin and C-peptide concentrations for the remaining 118 subjects. Seven subjects (one NGT, one IGT, and five T2DM subjects) were excluded because their model parameters were detected as outlier based on the adjusted outlyingness (Materials and Methods), and we analyzed the model for the remaining 111 subjects (47 NGT, 17 IGT, and 47 T2DM) (*Figure 2-source data 3*).

### Bidirectional changes of hepatic and peripheral insulin clearance during progression of glucose intolerance

We statistically compared the model parameters among the NGT, IGT, and T2DM groups (*Figure 2B* and Materials and Methods). Four of the nine parameters, *k*_*I*out_, *k*_ratio_, *h*, and *X*_b_, were significantly different.

The parameter *k*_*I*out_ is the degradation rate of serum insulin and corresponds to peripheral insulin clearance. *k*_*I*out_ in the NGT subjects was higher than that in the IGT and T2DM subjects (*Figure 2C*), indicating that peripheral clearance decreases during the progression of glucose intolerance, which is consistent with previous studies (Ohashi et al., 2015; Polidori et al., 2016).

The parameter *k*_ratio_ is the ratio of post-hepatic insulin to C-peptide, and (1 – *k*_ratio_) corresponds to the insulin extracted by the liver, that is, hepatic insulin clearance. (1 – *k*_ratio_) in the NGT subjects was lower than that in the IGT and T2DM subjects (*Figure 2C*), indicating the increase of hepatic insulin clearance in the IGT and T2DM subjects. This is consistent with an earlier clinical observation (Tamaki et al., 2013).

The parameter *h* is the threshold of plasma glucose concentration for the insulin secretion and corresponds to the fasting plasma glucose concentration. This parameter in the T2DM subjects was significantly higher than that in the NGT subjects (*Figure 2B*), consistent with the fact that fasting plasma glucose concentration is higher in T2DM (Abdul-Ghani et al., 2006; Hayashi et al., 2013).

The parameter *X*_b_ is the initial value of *X*, which may correspond to stored amounts of insulin and C-peptide or β-cell masses before the start of the hyperglycemic clamp. This parameter in the T2DM subjects was significantly lower than that in the NGT subjects (*Figure 2B*), consistent with observations that β-cell masses and stored insulin decrease in T2DM patients (Butler et al., 2003; Inaishi et al., 2016; Mizukami et al., 2014).

Using the same clamp data, we previously showed that insulin secretion decreases with the progression of glucose intolerance (Ohashi et al., 2015; Okuno et al., 2013). In this study, however, the parameters *α* and *β*, related to insulin secretion, did not show any significant differences among the NGT, IGT, and T2DM subjects, possibly because insulin secretion in this model also depends on other parameters such as *h, m, X*_b_, and *k*_ratio_, and the parameters involved in insulin secretion are too diverse.

The parameters *k*_ratio_, *k*_*I*out_, *k*_*CP*out_, *h*, and *X*_b_ show smaller variations than others (*Figure 2B*). This is probably because these parameters are directly related to the measured concentrations of serum insulin and C-peptide and plasma glucose, and therefore can be accurately estimated, whereas other parameters are not, resulting in large variation possibly due to inaccurate estimation.

### Relationship between hepatic and peripheral insulin clearance parameters and clinical indices of serum insulin regulation

We examined the correlation of the estimated model parameters with clinical indices of circulating insulin regulation among 111 subjects (*Figure 3; Figure 3-source data 1*). The model parameter showing the highest correlation with insulin sensitivity index (ISI) and with the metabolic clearance rate (MCR), which is the index of insulin clearance (see Materials and Methods for details), was peripheral insulin clearance, *k*_*I*out_ (*r* = 0.761 and 0.790, respectively, both *P* < 0.001). This correlation is consistent with our previous finding that peripheral insulin clearance is highly correlated with ISI and MCR (Ohashi et al., 2015). *k*_*I*out_ is the degradation rate of serum insulin, which depends on the number of insulin receptors on target tissues (Duckworth et al., 1998), indicating that serum insulin degradation and insulin sensitivity are mutually correlated. Therefore, it is reasonable that *k*_*I*out_ is correlated not only with MCR but also with ISI.

**Figure 3.**
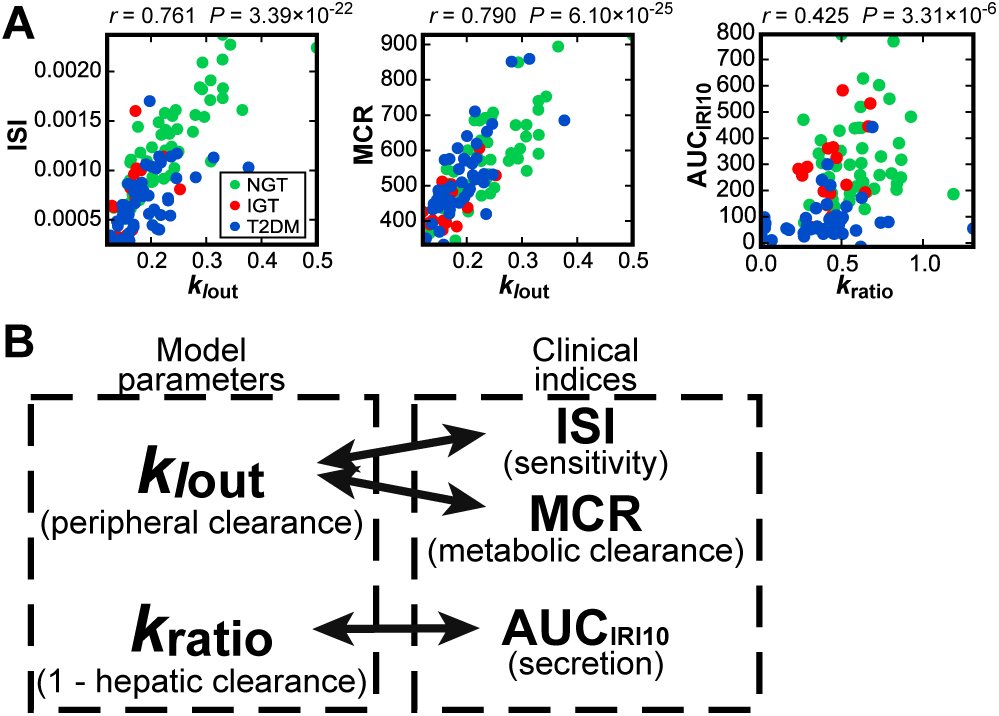
Model parameters showing the highest correlation with clinical indices. (A) Scatter plots for the indicated measured clinical indices versus the highest correlated model parameters (*Figure 3-source data 1*). ISI, the insulin sensitivity index; MCR, the metabolic clearance rate; AUC _IRI10_, amount of insulin secretion during the first 10 min of hyperglycemic clamp. Each dot indicates the values of an individual subject. The correlation coefficient, *r*, and the *P*-values for testing the hypothesis of no correlation are shown. (B) Summary of the model parameters *k*_*I*out_ and *k*_ratio_ showing the highest correlation with the indicated clinical indices.

The model parameter showing the highest correlation with insulin secretion during the first phase, AUC_IRI10_ (see Materials and Methods), which is the index of insulin secretion, was *k*_ratio_ (*r* = 0.425, *P* < 0.01). Note that (1 – *k*_ratio_) corresponds to hepatic insulin clearance. Because the parameter *k*_ratio_ is the fraction of insulin remaining after the hepatic extraction, its correlation with insulin secretion is reasonable.

In addition, the model parameter showing the highest correlation with both FPG and 2-h PG, the main indices of glucose tolerance, was *h* (*r* = 0.448 and 0.504, respectively, both *P* < 0.001), which is the threshold glucose concentration for insulin secretion. This finding is consistent with *h* corresponding to fasting plasma glucose concentration. The model parameter showing the highest correlation with the clamp disposition index, clamp DI, which is calculated as the product of insulin secretion AUC_IRI10_ and ISI and is the index of glucose tolerance (Okuno et al., 2013), was *k*_ratio_ · *k*_*I*out_ (*r* = 0.540, *P* < 0.001). Considering that *k*_ratio_ is related to post-hepatic insulin delivery, and *k*_*I*out_ is related to insulin sensitivity, which depends on the number of insulin receptors on target organs, it is reasonable that the product *k*_ratio_ · *k*_*I*out_ shows the highest correlation with clamp DI, which is also the product of clinically estimated insulin secretion and sensitivity.

### Selective regulation of amplitude and temporal patterns of serum insulin concentration by hepatic and peripheral insulin clearance

Because *k*_*I*out_ and *k*_ratio_ were the parameters showing the highest correlation with clinical indices of insulin sensitivity and secretion, respectively, both of which are related to the progression of glucose intolerance and T2DM, we analyzed the roles of *k*_ratio_ and *k*_*I*out_ in the temporal changes of serum insulin concentration (*Figure 4*). We changed the originally estimated values of *k*_ratio_ or *k*_*I*out_ or both by 2^−1^ to 2^1^ times and simulated the time course of *I*, serum insulin concentration, during hyperglycemic clamp for each subject (*Figure 4-figure supplement 1*). Similar temporal changes of *I* versus changes in the parameters were observed in all 111 subjects, so only the simulation result of subject #3 (NGT) is shown (*Figure 4A*).

**Figure 4.**
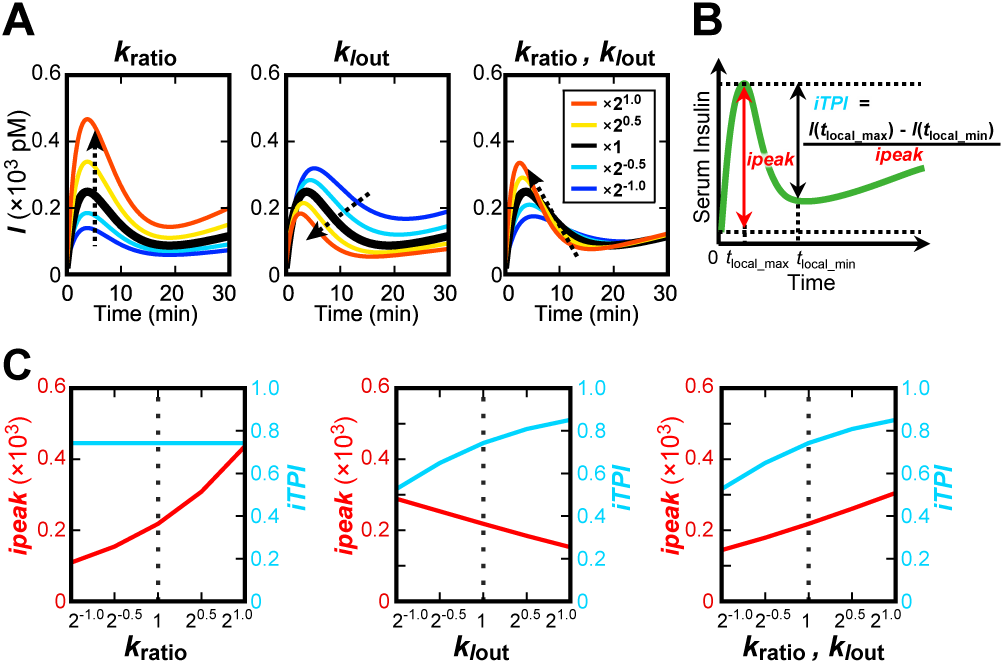
The roles of *k*_ratio_ and *k*_*I*out_ in the amplitude and temporal patterns of serum insulin concentration. (A) Simulated time course of serum insulin concentration *I* during hyperglycemic clamp of subject #3 by changing *k*_ratio_ or *k*_*I*out_ or both as indicated (see Materials and Methods). Dotted arrows indicate the direction of the change in the temporal pattern as the parameter increases. (B) The definition of *ipeak* (incremental peak) and *iTPI* (incremental transient peak index), reflecting the peak amplitude and the temporal pattern of serum insulin concentration *I*. (C) *ipeak* and *iTPI* of *I* of subject #3 by changing *k*_ratio_ or *k*_*I*out_ or both.

The time course of *I* with the original parameters in the model of subject #3 showed the transient increase (*Figure 4A*, black line). As *k*_ratio_ increased, *I* increased without changing the transient pattern (*Figure 4A*, left panel, red line). Indeed, an increase of *k*_ratio_ affects the value of *I* similarly at any time point, because *k*_ratio_ controls the gain of time derivative of *I*. As *k*_*I*out_ increased, *I* decreased and the temporal pattern became more transient with an earlier peak time (*Figure 4A*, middle panel, red line). Conversely, as *k*_*I*out_ decreased, *I* increased and the temporal pattern became more sustained with a delayed peak time (*Figure 4A*, middle panel, blue line). This result suggests that *k*_*I*out_ controls the shift in the temporal patterns of *I* from transient to sustained. These changes in the temporal pattern of *I* are characterized by a relative decrease in the first-phase secretion and relative increase in the second-phase secretion. Note that the decrease in *k*_*I*out_ also increases the amplitude of *I*.

Because both *k*_ratio_ and *k*_*I*out_ decreased during the progression of glucose intolerance (*Figure 2B*), we examined the effect of the simultaneous changes of *k*_ratio_ and *k*_*I*out_ on the amplitude and transient/sustained patterns of *I*. When both *k*_ratio_ and *k*_*I*out_ increased with the same ratio, *I* increased during first-phase secretion (0–10 min), whereas *I* decreased during second-phase secretion (10–30 min) (*Figure 4A*, right panel, red line). Thus, simultaneous increase of *k*_ratio_ and *k*_*I*out_ results in the increase of peak amplitude of *I* and in changes in the temporal pattern of *I* from sustained to transient.

We quantified the role of *k*_ratio_ and *k*_*I*out_ in the peak amplitude and temporal patterns of *I*. We defined the index *ipeak* (incremental peak) for the peak amplitude of *I*, and the index *iTPI* (incremental transient peak index; modified from Kubota et al., 2012) for the temporal pattern of *I* (*Figure 4B*), as follows:

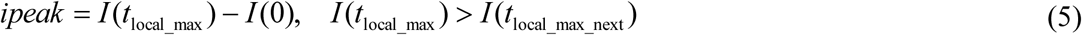

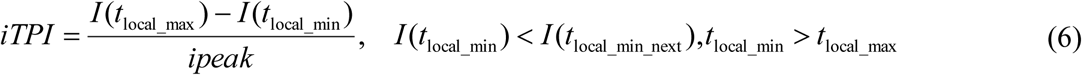
where *I*(*t*) represents *I* at time *t, t*_local_max_ is the time at which *I* stops increasing for the first time from 0 min, *t*_local_max_next_ is the next sampling time of *t*_local_max_, *t*_local_min_ is the time at which *I* stops decreasing for the first time after *t*_local_max_, and *t*_local_min_next_ is the next sampling time of *t*_local_min_.

The index *ipeak* is the difference in *I* between the local maximum *I*(*t*_local_max_) and the initial fasting concentration *I*(0) and represents the peak amplitude of *I* during the first-phase secretion. The index *iTPI* is the ratio of the difference of *I* between the local maximum *I*(*t*_local_max_) and the local minimum *I*(*t*_local_min_) of *I* against *ipeak*, which reflects the ratio of *I* during the first- and second-phase secretions. As *iTPI* approaches 1, the difference in *I* between the first- and second-phase secretions becomes larger, meaning that the temporal change of *I* becomes more transient. Conversely, as *iTPI* approaches 0, the difference in *I* between the first- and second-phase secretions becomes smaller, meaning that the temporal change of *I* becomes more sustained.

We calculated *ipeak* and *iTPI* from the simulated time courses of *I* by changing the original estimates of *k*_ratio_ or *k*_*I*out_ or both by 2^−1^ to 2^1^ times. As *k*_ratio_ increased, *ipeak* increased but *iTPI* did not change (*Figure 4C*, left panel), indicating that increasing *k*_ratio_ increases the peak amplitude of *I* during the first-phase secretion without changing its temporal pattern. As *k*_*I*out_ increased, *iTPI* increased and *ipeak* decreased (*Figure 4C*, middle panel), indicating that increasing *k*_*I*out_ changes the temporal patterns of *I* from sustained to transient and decreases the peak amplitude of *I* during the first-phase secretion.

When both *k*_ratio_ and *k_Iout_* increased at the same ratio, both *ipeak* and *iTPI* increased (*Figure 4C*, right panel), indicating that increasing both *k*_ratio_ and *k*_*I*out_ increases the peak amplitude of *I* and changes the temporal pattern from sustained to transient. The increase in *ipeak* means that the effect of *k*_ratio_, which increases *ipeak*, is stronger than that of *k_Iout_*, which decreases *ipeak*. Given that both *k*_ratio_ and *k_Iout_* decrease during the progression of glucose intolerance, both *ipeak* and *iTPI* decrease (*Figure 5*). This finding is consistent with earlier clinical observations that the peak amplitude of circulating insulin concentration during the first-phase secretion decreases and the temporal pattern becomes more sustained during the progression of glucose intolerance (Cerasi and Luft, 1967; Del Prato and Tiengo, 2001; Seino et al., 2011).

**Figure 5.**
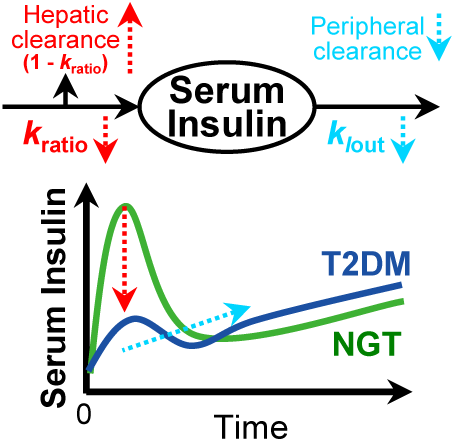
Bidirectional changes of hepatic and peripheral insulin clearance during the progression of glucose intolerance. During the progression from NGT to T2DM, hepatic insulin clearance (1 – *k*_ratio_) increases and peripheral insulin clearance *k*_*I*out_ decreases, characterizing the decrease in peak amplitude and the change in the temporal pattern of serum insulin concentration from transient to sustained, respectively.

We performed parameter sensitivity analysis on *ipeak* and *iTPI* in the simulation for each model parameter (*Table 2*, Materials and Methods). We compared the median of the parameter for all 111 subjects as the parameter sensitivity index (*Table 2*). For *ipeak* the *k*_ratio_ had the significantly highest median, and for *iTPI k_Iout_* showed the significantly highest median, indicating that hepatic insulin clearance and peripheral insulin clearance are the most critical parameters controlling the peak amplitude and temporal patterns of serum insulin concentration, respectively.

**Table 2.** Parameter sensitivity analysis for *ipeak* and *iTPI*.

| Rank | *ipeak* |  |  | *iTPI* |  |  |
| --- | --- | --- | --- | --- | --- | --- |
|  |  | Median | *P* |  | Median | *P* |
| 1 | $k_{\text{ratio}}$ | 1.00 | 1.00 | $k_{\text{Iout}}$ | $4.64 \times 10^{-1}$ | 1.00 |
| 2 | $m$ | $3.52 \times 10^{-1}$ | $6.26 \times 10^{-38}$ | $\beta$ | $-2.53 \times 10^{-1}$ | $1.30 \times 10^{-6}$ |
| 3 | $k_{\text{Iout}}$ | $-2.74 \times 10^{-1}$ | $1.23 \times 10^{-36}$ | $\alpha$ | $-1.42 \times 10^{-1}$ | $5.36 \times 10^{-15}$ |
| 4 | $h$ | $-3.64 \times 10^{-4}$ | $4.04 \times 10^{-34}$ | $h$ | $1.38 \times 10^{-1}$ | $8.70 \times 10^{-7}$ |
| 5 | $\beta$ | $2.59 \times 10^{-4}$ | $1.23 \times 10^{-36}$ | $m$ | $7.18 \times 10^{-2}$ | $2.56 \times 10^{-9}$ |
| 6 | $\alpha$ | $1.61 \times 10^{-4}$ | $1.23 \times 10^{-36}$ | $k_{\text{ratio}}$ | $8.28 \times 10^{-15}$ | $2.62 \times 10^{-35}$ |

Medians of the indicated parameters for all 111 subjects were used for parameter sensitivity analysis for *ipeak* and *iTPI* (see Materials and Methods). The higher the absolute value of the parameter, the higher the sensitivity. *P*-values relative to the median for the top-ranked parameter were determined by two-sided Wilcoxon rank sum test, and the value of $<4.17 \times 10^{-3}$ ( $=0.05/12$ , corrected by the number of tests, divided by 12) is considered statistically significant.

The measured temporal changes of the serum insulin concentration during hyperglycemic clamp (0–90 min) showed no clear difference between the NGT and IGT subjects (*Figure 1*). However, the estimated *k*_ratio_ and *k_Iout_* in the NGT subjects were significantly higher than those in the IGT subjects, suggesting that hepatic and peripheral insulin clearance are not the only parameters responsible for the peak amplitude and temporal patterns of serum insulin concentration, respectively. Other parameters such as insulin secretion, which also affects the temporal changes of serum insulin concentration, may compensate for the temporal changes by insulin clearance between the NGT and IGT subjects. This also means that changes in the hepatic and peripheral insulin clearance during the progression of glucose intolerance cannot be directly assessed from the measured time course of serum insulin concentration, but must be evaluated with a mathematical model.

From Eq. 3, the parameter *k*_ratio_ directly reflects post-hepatic insulin delivery and is involved in the increase in gain of *I*. Therefore, the change of *k*_ratio_ is directly reflected in the peak amplitude, so the sensitivity to *ipeak* becomes 1. This means that hepatic insulin clearance is more responsible for the peak amplitude of serum insulin concentration than the pre-hepatic insulin secretion before extraction by the liver.

The parameter *k_Iout_* corresponds to peripheral insulin clearance and is the degradation rate of *I*, meaning that *k_Iout_* is the time constant of serum insulin degradation. *k_Iout_* is the parameter directly responsible for temporal conversion of input *X*, delivered insulin after the extraction by the liver, into output, *I*. Thus, *k_Iout_* is the most sensitive parameter for *iTPI* corresponding to the shift between the transient and sustained temporal pattern of serum insulin concentration.

## Discussion

We developed several alternative mathematical models using concentrations of plasma glucose and serum insulin and C-peptide during consecutive hyperglycemic and hyperinsulinemic-euglycemic clamps, and selected the model showing the best fit for all subjects. The fact that a single model with the same structure and different parameters was selected for all subject means that the progression of glucose intolerance from NGT to IGT to T2DM is not qualitative, but quantitative and continuous.

During the progression of glucose intolerance, it has been shown that the peak amplitude of circulating insulin concentration during the first-phase secretion decreases and the temporal pattern becomes more sustained (Cerasi and Luft, 1967; Del Prato and Tiengo, 2001; Seino et al., 2011). In this study, we found that both *k*_*I*out_, corresponding to peripheral insulin clearance, and *k*_ratio_ decrease as glucose intolerance progressed. Given that (1 – *k*_ratio_), corresponding to hepatic insulin clearance, increases as the *k*_ratio_ decreases, our finding strongly suggests that, during the progression of glucose intolerance, the peak amplitude of serum insulin concentration decreases due to the increase in hepatic insulin clearance and the temporal pattern changes from transient to sustained occur because of a decrease in peripheral insulin clearance (*Figure 5*). Importantly, the decrease in peripheral insulin clearance alone can explain only the temporal change of serum insulin concentration, not the decrease of peak amplitude. Thus, the increase of hepatic insulin clearance and decrease of peripheral insulin clearance simultaneously cause the decrease in the peak amplitude of serum insulin concentration during the first-phase secretion and change in temporal pattern from transient to sustained. Our result demonstrates that, in addition to the decrease in insulin secretion (Del Prato and Tiengo, 2001; Seino et al., 2011), the increase in hepatic clearance also contributes to the decrease in peak amplitude of serum insulin concentration in the first-phase secretion during the progression of glucose intolerance.

In our model, as *k*_ratio_, the ratio of post-hepatic insulin compared to C-peptide, increased, *ipeak*, the peak amplitude of peripheral insulin concentration, increased (*Figure 4C*). According to clinical measurements, *k*_ratio_ was also correlated with *ipeak* calculated directly from the serum insulin concentration measured during the first-phase secretion in hyperglycemic clamp (*Figure 3-figure supplement 1B, r* = 0.423, *P* < 0.001), indicating that hepatic insulin clearance is pathologically correlated with the peak amplitude in the first-phase secretion during the progression of glucose intolerance. On the other hand, in our model, as *k*_*I*out_, the peripheral insulin clearance, increased, *iTPI*, representing the temporal pattern of peripheral insulin concentration, increased (*Figure 4C*). However, in clinical measurements, *k*_*I*out_ was not highly correlated with *iTPI* calculated directly from the serum insulin concentration measured during hyperglycemic clamp (*Figure 3-figure supplement 1B, r* = 0.297, *P* < 0.01). The reason for this lack of correlation between *k_Iout_* and *iTPI* in clinical measurement remains unclear; however, it may be because little insulin was secreted during hyperglycemic clamp in some IGT and T2DM subjects, and *iTPI* cannot be estimated accurately because of low concentration of serum insulin.

Many studies have shown that insulin clearance decreases in T2DM patients (Benzi et al., 1994; Jones et al., 1997; Lee et al., 2013; Marini et al., 2014; Polonsky et al., 1988a; Rudovich et al., 2004). However, the change in hepatic insulin clearance in this condition has been controversial, with some studies finding an increase in T2DM subjects (Tamaki et al., 2013) and others a decrease (Bonora et al., 1983; Marini et al., 2014). We previously developed a mathematical model using data gathered during hyperglycemic and hyperinsulinemic-euglycemic clamps, and peripheral insulin clearance significantly decreased during the progression toward T2DM (Ohashi et al., 2015). However, hepatic and peripheral insulin clearances were not estimated separately because we did not use C-peptide data. Recently, Polidori et al. (2016) estimated both hepatic and peripheral insulin clearance by modeling analysis using plasma insulin and C-peptide concentrations obtained from the insulin-modified frequently sampled IVGTT. They found that the peripheral insulin clearance significantly decreased in IGT subjects compared with NGT subjects, whereas hepatic insulin clearance did not significantly differ between the IGT and NGT subjects (Polidori et al., 2016); the former finding is consistent with our result that peripheral insulin clearance decreases during the progression of glucose intolerance. We demonstrated that hepatic insulin clearance significantly increases, whereas peripheral insulin clearance significantly decreases during the progression from NGT to T2DM. The increase in hepatic insulin clearance may be caused by impaired suppression of endocytosis of insulin receptors on the liver (Tamaki et al., 2013), and the decrease in peripheral insulin clearance may be caused by a decrease of the number of insulin receptors on target tissues (Duckworth et al., 1998). Polidori et al. (2016) also found that hepatic and peripheral insulin clearances were not highly correlated. Consistent with their results, in our analysis the insulin clearance parameters *k*_ratio_ and *k*_*I*out_ were not highly correlated (*Figure 3-figure supplement 1A, r* = 0.296, *P* < 0.01), suggesting that both insulin clearances are independently regulated.

Insulin selectively regulates various functions, such as signaling activities, metabolic control, and gene expression, depending on its temporal patterns. For example, we previously reported that pulse stimulation of insulin in rat hepatoma Fao cells, resembling the first-phase secretion, selectively regulated glycogen synthase kinase-3β (GSK3β), which regulates glycogenesis, and S6 kinase, which regulates protein synthesis, whereas ramp stimulation of insulin, resembling the second-phase secretion, selectively regulated GSK3β and glucose-6-phosphatase (*G6Pase*),which regulates gluconeogenesis (Kubota et al., 2012). We also found that insulin-dependent metabolic control and gene expression are selectively regulated by temporal patterns and doses of insulin in FAO cells (Noguchi et al., 2013; Sano et al., 2016). Sustained stimulation of insulin suppressed the expression of insulin receptors, leading to reduced insulin sensitivity in FAO cells (Goodner et al., 1988; Hirashima et al., 2003; Hori et al., 2006). Likewise, phosphorylation of the insulin receptor substrate (IRS)-1/2 in rat liver increased when pulsatile (rather than continuous) stimulation of insulin was imposed in the portal circulation (Matveyenko et al., 2012). This may have occurred through the negative feedback within the insulin signaling pathway, the phosphatidylinositide (PI) 3-kinase/Alt pathway, targeting IRS-1/2 (Hirashima et al., 2003; Sedaghat et al., 2002). In addition, IRS-2, rather than IRS-1, mainly regulates hepatic gluconeogenesis through its rapid downregulation by insulin (Kubota et al., 2008), suggesting the selective roles of IRS-1/2 in response to temporal patterns of plasma insulin. These findings indicate that the amplitude and temporal pattern of circulating insulin concentration selectively regulate insulin actions on the target tissues. Given that hepatic and peripheral insulin clearances are responsible for the amplitude and temporal pattern of circulating insulin concentration, these clearances are likely to be involved in selective control of insulin action, glucose homeostasis, and the pathogenesis of T2DM.

We previously developed a mathematical model for concentrations of plasma glucose and serum insulin measured during consecutive hyperglycemic and hyperinsulinemic-euglycemic clamps and found significant decreases in insulin secretion, sensitivity, and peripheral insulin clearance during the progression from NGT to IGT to T2DM (Ohashi et al., 2015). The differences between our previous study and this study are the model structure and C-peptide data. The previous model consisted of plasma glucose and serum insulin and required only glucose and insulin infusion as inputs. The model in this study does not have plasma glucose concentration but includes serum insulin and C-peptide concentrations, while plasma glucose concentration and insulin infusion are used as inputs (*Figure 2A*; *Figure 2-figure supplement 1*). In the previous study, only peripheral insulin clearance, but not hepatic insulin clearance, was estimated because C-peptide data were not used. The decrease of insulin clearance from NGT to T2DM in the previous study is consistent with the decrease of peripheral insulin clearance from NGT to T2DM in this study. In the previous study, the parameter corresponding to insulin secretion in the NGT and IGT subjects was significantly higher than that in the T2DM subjects; however, the parameter related to insulin secretion did not show a significant difference between the NGT, IGT, and T2DM subjects in this study, possibly because the parameters related to insulin secretion (*α, β, h, m, X*_b_, and *k*_ratio_) are too diversified. The parameter corresponding to insulin sensitivity was not incorporated in this study.

Many mathematical models to reproduce circulating C-peptide concentration have been developed. A two-compartment model for C-peptide kinetics was originally proposed (Eaton et al., 1980). A combined model that included both circulating insulin and C-peptide kinetics described by a single compartment structure was introduced to estimate hepatic insulin clearance (Vølund et al., 1987). The C-peptide minimal model describing peripheral insulin and C-peptide appearance and kinetics was also developed to assess hepatic insulin clearance (Cobelli and Pacini, 1988; Grodsky, 1972; Ličko and Silvers, 1975; Toffolo et al., 2001; Toffolo et al., 2006), and several other model structures for circulating C-peptide concentration were reported (Chase et al., 2010; Mari et al., 2002). One difference between others’ and our studies is the experimental protocol in which data were applied to parameter estimation. IVGTT or hyperglycemic clamp were performed for parameter estimation in models of circulating C-peptide concentration, whereas we used hyperglycemic and hyperinsulinemic-euglycemic clamps, which may improve the accuracy of the parameter estimation of peripheral insulin clearance, *k*_*I*out_. Recently, a model of plasma insulin concentration including hepatic and peripheral insulin clearance and the delivery of insulin from the systemic circulation to the liver during the insulin-modified IVGTT was proposed (Polidori et al., 2016). In that model, the parameter of hepatic insulin clearance was negatively correlated with acute insulin secretion in response to glucose, and the parameter of peripheral insulin clearance was correlated with insulin sensitivity (Polidori et al., 2016), consistent with the results in this study. The high correlation between the parameter of hepatic insulin clearance, *k_ratio_*, and the clinical index of insulin secretion, AUC_IRI10_, suggests the possibility that hepatic insulin clearance considerablely affects the clinical index of insulin secretion measured by peripheral insulin concentration because the clinical index of insulin secretion, AUC_IRI10_, was measured by the post-hepatic insulin delivery, and therefore reflects both insulin secretion and hepatic insulin clearance. This suggests that insulin secretion *per se* in the clinical index of insulin secretion may be overestimated because of the involvement of hepatic insulin clearance. Further study is neccesary to address this issue.

In conclusion, using the mathematical model for serum insulin and C-peptide concentrations during consecutive hyperglycemic and hyperinsulinemic-euglycemic clamps, we determined the quantitative structure of the control of circulating insulin concentration. The estimated model parameters revealed the increase of hepatic insulin clearance and decrease of peripheral insulin clearance during the progression of T2DM, and these changes selectively regulate the amplitude and temporal patterns of serum insulin concentration, respectively. The bidirectional changes of both types of clearance shed light on the pathological mechanism underlying the abnormal temporal patterns of circulating insulin concentration during the progression of glucose intolerance.

## Materials and Methods

### Subjects and measurements

The plasma and serum measurement data originated from our previous research (Ohashi et al., 2015; Okuno et al., 2013). This metaboic analysis was approved by the ethics committee of Kobe University Hospital, and written informed consent was obtained from all subjects. In brief, 50 NGT, 18 IGT, and 53 T2DM subjects underwent the consecutive clamp analyses. From 0 to 90 min, a hyperglycemic clamp was applied by intravenous infusion of a bolus of glucose (9622 mg/m^2^) within 15 min followed by that of a variable amount of glucose to maintain the plasma glucose level at 200 mg/dL. Ten minutes after the end of the hyperglycemic clamp, a 120-min hyperinsulinemic-euglycemic clamp was initiated by intravenous infusion of human regular insulin (Humulin R, Eli Lilly Japan K.K.) at a rate of 40 mU m^−2^ min^−1^ and with a target plasma glucose level of 90 mg/dL. For the NGT and IGT subjects whose plasma glucose levels were <90 mg/dL, the plasma glucose concentration was clamped at the fasting level. We measured the plasma glucose level every 1 min during the clamp analyses and obtained the 5-min average values. We also measured insulin and C-peptide level in serum samples collected at 5, 10, 15, 60, 75, 90, 100, 190, and 220 min after the onset of the tests. First-phase insulin secretion during the hyperglycemic clamp was defined as the incremental area under the immunoreactive insulin (IRI) concentration curve (μU mL^−1^ min^−1^) from 0 to 10 min (AUC_IRI10_). The insulin sensitivity index (ISI) derived from the hyperinsulinemic-euglycemic clamp was calculated by dividing the mean glucose infusion rate during the final 30 min of the clamp (mg kg^−1^ min^−1^) by both the plasma glucose (mg/dL) and serum insulin (μU/mL) levels at the end of the clamp and then multiplying the result by 100. A clamp-based analog of the disposition index, the clamp disposition index (clamp DI), was calculated as the product of AUC_IRI10_ and ISI, as described previously (Okuno et al., 2013). The metabolic clearance rate (MCR) (DeFronzo et al., 1979), an index of insulin clearance, was calculated by dividing the insulin infusion rate at the steady state (1.46 mU kg^−1^ min^−1^) by the increase in insulin concentration above the basal level in the hyperinsulinemic-euglycemic clamp (Okuno et al., 2013): 1.46 (mU kg^−1^ min^−1^) × body weight (kg) × body surface area (m^2^)/(end IRI – fasting IRI) (μU/mL), where body surface area is defined as (body weight (kg))^1/2^ × (body height (cm))^1/2^ / 60 (Mosteller formula). The actual data for all 121 subjects are shown in *Figure 1-source data 1*.

### Mathematical models

We developed six mathematical models based on the proposed models in order to choose the best model for reproducing our measurement of serum insulin and C-peptide during consecutive hyperglycemic and hyperinsulinemic-euglycemic clamps (*Figure 2-figure supplement 1*). In these models, *I* represents serum insulin concentration (pM), and *CP* and *CP*_1_ represent serum C-peptide concentration (pM) including insulin and C-peptide secretion and hepatic and peripheral clearance. We used a conversion factor of insulin (6.00 nmol/U) (Vølund et al., 1991) and the molecular weights of glucose (180.16) and C-peptide (3020.3) to convert the unit of serum insulin, plasma glucose, and serum C-peptide, respectively. We used plasma glucose concentration *G* (mM) as input in the models, which was determined by linear interpolation of the measured plasma glucose data. Note that plasma glucose data were obtained as the 5-min average values, and each sampling time was reduced by 2 min in the calculation of linear interpolation.

The actual insulin infusion rate (*IIR*, mU kg^−1^ min^−1^) was converted to the corresponding serum concentrations (*cIIR*) as follows:

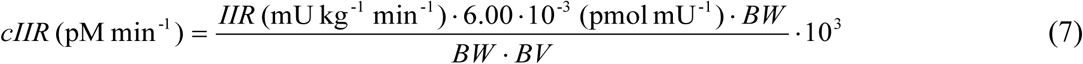
where *BW* and *BV* denote body weight and blood volume (75 and 65 mL/kg for men and women, respectively, Choi and Ahn, 2011).

In the models, insulin infusions are represented by *influx*. This flux follows the nonlinear function *f* that predicts insulin infusion concentrations. Given that insulin infusion was performed only during the hyperinsulinemic-euglycemic (from 100 to 220 min) clamp, the function *f* was given by the following equations:

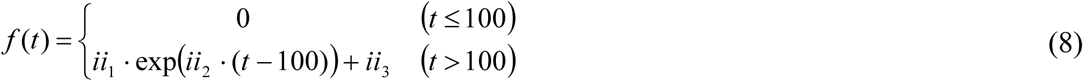
where the parameters *ii_j_* (*j* = 1, 2, 3) are estimated to reproduce *cIIR* for each subject with a nonlinear least squares technique (Lagarias et al., 1998).

### Parameter estimation

The model parameters for each subject were estimated to reproduce the experimentally measured time course by a meta-evolutionary programming method to approach the neighborhood of the local minimum, followed by application of the nonlinear least squares technique to reach the local minimum (Fujita et al., 2010). Each parameter was estimated in the range from 10^−6^ to 10^6^. For these methods, the model parameters were estimated to minimize the objective function value, which is defined as the residual sum of squares (RSS) between the actual time course obtained by clamp analyses and the model trajectories. *RSS* is given by:

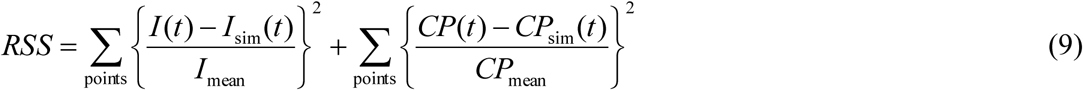
where 
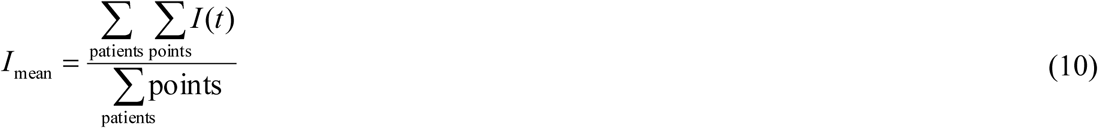

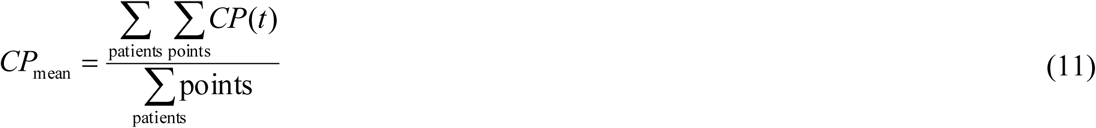
*I*(*t*) and *CP*(*t*) are the serum insulin and C-peptide concentration, and *I*_sim_(*t*) and *CP*_sim_(*t*) are simulated serum insulin and C-peptide concentrations at *t* min, respectively. Serum insulin and C-peptide concentrations were normalized by dividing them by the averages of serum concentrations over all time points of all subjects of insulin (*I*_mean_, 302.7 pM) and C-peptide (*CP*_mean_, 1475 pM), respectively. The numbers of parents and generations in the meta-evolutionary programming were 400 and 4000, respectively. Model parameters for all subjects are shown in *Figure 2-source data 1*.

### Model selection

The best model to reproduce serum insulin and C-peptide time courses with a simpler model structure among the six models was chosen according to Akaike information criterion (AIC). For a given model and a single subject, AIC was calculated as follows:

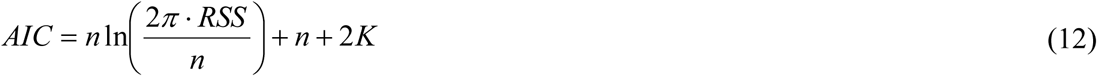
where *n* is the total number of sampling time points of serum insulin and C-peptide, and *K* is the number of estimated parameters of the model.

### Determination of parameter outliers

The outliers of RSS and model parameters were detected by the adjusted outlyingness (AO) (Hubert and Van der Veeken, 2008). The cutoff value of AO was *Q_3_* + 1.5e*^3MC^* · *IQR*, where *Q_3_, MC*, and *IQR* are the third quartile, medcouple, and interquartile range, respectively. The medcouple is a robust measure of skewness (Brys et al., 2004). The number of directions was set at 8000. Subjects found to have outlier parameters (1 NGT, 1 IGT, and 5 T2DM subjects) were excluded from further study.

### Parameter sensitivity analysis

We defined the individual model parameter sensitivity for each subject as follows:

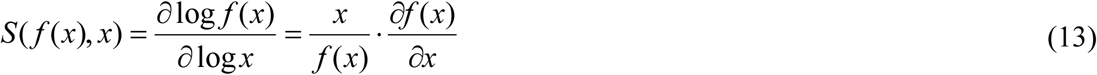
where *x* is the parameter value and *f*(*x*) is *ipeak* or *iTPI*. The differentiation is numerically approximated by central difference 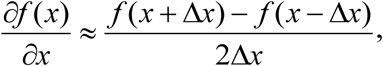, and *x* + Δ*x* and *x* – Δ*x* were set so as to be increased [*x* (1.1*x*)] or decreased [*x* (0.9*x*)] by 10%, respectively. Finally, we defined the parameter sensitivity by the median of the individual parameter sensitivity for all subjects. We examined the parameter sensitivity for six parameters of the rate constant related to serum insulin concentration, except *X_b_* and *k*_*CP*out_. The higher the absolute value of parameter sensitivity, the larger the effect of the parameter on *ipeak* or *iTPI*.

### Statistical analysis

Unless indicated otherwise, data are expressed as the median with first and third quartiles. Medians of parameter values were compared among the NGT, IGT, and T2DM subjects with the use of the Steel-Dwass test (Neuhauser and Bretz, 2001), a statistical nonparametric test for multiple comparisons. A *P* value <0.05 was considered statistically significant.

## Acknowledgements

We thank fellow laboratory members for critical reading of the manuscript and for technical assistance with the analysis. The computations for this work were performed in part on the NIG supercomputer system at ROIS National Institute of Genetics. This work was supported by the Creation of Fundamental Technologies for Understanding and Control of Biosystem Dynamics, CREST, of the Japan Science and Technology Agency (JST); Japan Diabetes Foundation; and Ono Medical Research Foundation

## Author contributions

K.O., M.F., W.O., and S.K. conceived the project; H. Komada, K.S., and W.O. designed and performed the experiments; K.O. and M.F. developed the computational model; K.O., M.F., S.U., and H. Kubota analyzed the data; K.O., W.O. and S.K. wrote the manuscript.

## References

Abdul-Ghani, M. A., Tripathy, D. & Defronzo, R. A. 2006. Contributions of beta-cell dysfunction and insulin resistance to the pathogenesis of impaired glucose tolerance and impaired fasting glucose. Diabetes Care, 29, 1130–9.

AhrÉN, B. & Thorsson, O. 2003. Increased insulin sensitivity is associated with reduced insulin and glucagon secretion and increased insulin clearance in man. J Clin Endocrinol Metab, 88, 1264–70.

Akaike, H. 1973. Information theory and an extension of the maximum likelihood principle. In: Petrov, B. N. & CsÁKi, F. (eds.) Proceedings of the 2nd International Symposium on Information Theory. Budapest: Akadémiai Kiadó.

Alberti, K. G. & Zimmet, P. Z. 1998. Definition, diagnosis and classification of diabetes mellitus and its complications. Part 1. Diagnosis and classification of diabetes mellitus provisional report of a WHO consultation. Diabet Med, 15, 539–53.

Benzi, L., Cecchetti, P., Ciccarone, A., Pilo, A., Di Cianni, G. & Navalesi, R. 1994. Insulin degradation in vitro and in vivo: a comparative study in men. Evidence that immunoprecipitable, partially rebindable degradation products are released from cells and circulate in blood. Diabetes, 43, 297–304.

Bergman, R. N., Phillips, L. S. & Cobelli, C. 1981. Physiologic evaluation of factors controlling glucose tolerance in man: measurement of insulin sensitivity and beta-cell glucose sensitivity from the response to intravenous glucose. J Clin Invest, 68, 1456–67.

Bonora, E., Zavaroni, I., Coscelli, C. & Butturini, U. 1983. Decreased hepatic insulin extraction in subjects with mild glucose intolerance. Metabolism, 32, 438–46.

Breda, E., Cavaghan, M. K., Toffolo, G., Polonsky, K. S. & Cobelli, C. 2001. Oral glucose tolerance test minimal model indexes of beta-cell function and insulin sensitivity. Diabetes, 50, 150–8.

Brys, G., Hubert, M. & Struyf, A. 2004. A robust measure of skewness. J Computat Graph Stat, 13, 996–1017.

Butler, A. E., Janson, J., Bonner-Weir, S., Ritzel, R., Rizza, R. A. & Butler, P. C. 2003. Beta-cell deficit and increased beta-cell apoptosis in humans with type 2 diabetes. Diabetes, 52, 102–10.

Cerasi, E. & Luft, R. 1967. The plasma insulin response to glucose infusion in healthy subjects and in diabetes mellitus. Acta Endocrinol (Copenh), 55, 278–304.

Chase, J. G., Mayntzhusen, K., Docherty, P. D., Andreassen, S., Mcauley, K. A., Lotz, T. F. & Hann, C. E. 2010. A three-compartment model of the C-peptide-insulin dynamic during the DIST test. Math Biosci, 228, 136–46.

Choi, E. & Ahn, W. 2011. A new mixture ratio of heparin for the cell salvage device. Korean J Anesthesiol, 60, 226.

Cobelli, C. & Pacini, G. 1988. Insulin secretion and hepatic extraction in humans by minimal modeling of C-peptide and insulin kinetics. Diabetes, 37, 223–31.

Cobelli, C., Dalla Man, C., Toffolo, G., Basu, R., Vella, A. & Rizza, R. 2014. The oral minimal model method. Diabetes, 63, 1203–13.

Defronzo, R. A., Tobin, J. D. & Andres, R. 1979. Glucose clamp technique: a method for quantifying insulin secretion and resistance. Am J Physiol, 237, E214–23.

Defronzo, R. A., Bonadonna, R. C. & Ferrannini, E. 1992. Pathogenesis of NIDDM: a balanced overview. Diabetes Care, 15, 318–68.

Del Prato, S. & Tiengo, A. 2001. The importance of first-phase insulin secretion: implications for the therapy of type 2 diabetes mellitus. Diabetes Metab Res Rev, 17, 164–74.

Duckworth, W. C., Hamel, F. G. & Peavy, D. E. 1988. Hepatic metabolism of insulin. Am J Med, 85, 71–6.

Duckworth, W. C., Bennett, R. G. & Hamel, F. G. 1998. Insulin degradation: progress and potential. Endocr Rev, 19, 608–24.

Eaton, R. P., Allen, R. C., Schade, D. S., Erickson, K. M. & Standefer, J. 1980. Prehepatic insulin production in man: kinetic analysis using peripheral connecting peptide behavior. J Clin Endocrinol Metab, 51, 520–8.

Fujita, K. A., Toyoshima, Y., Uda, S., Ozaki, Y., Kubota, H. & Kuroda, S. 2010. Decoupling of receptor and downstream signals in the Akt pathway by its low-pass filter characteristics. Sci Signal, 3, ra56.

Goodner, C. J., Sweet, I. R. & Harrison, H. C. 1988. Rapid reduction and return of surface insulin receptors after exposure to brief pulses of insulin in perifused rat hepatocytes. Diabetes, 37, 1316–23.

Grodsky, G. M. 1972. A threshold distribution hypothesis for packet storage of insulin and its mathematical modeling. J Clin Invest, 51, 2047–59.

Hayashi, T., Boyko, E. J., Sato, K. K., Mcneely, M. J., Leonetti, D. L., Kahn, S. E. & Fujimoto, W. Y. 2013. Patterns of insulin concentration during the OGTT predict the risk of type 2 diabetes in Japanese Americans. Diabetes Care, 36, 1229–35.

Hirashima, Y., Tsuruzoe, K., Kodama, S., Igata, M., Toyonaga, T., Ueki, K., Kahn, C. R. & Araki, E. 2003. Insulin down-regulates insulin receptor substrate-2 expression through the phosphatidylinositol 3-kinase/Akt pathway. J Endocrinol, 179, 253–66.

Hori, S. S., Kurland, I. J. & Distefano, J. J. 2006. Role of endosomal trafficking dynamics on the regulation of hepatic insulin receptor activity: models for Fao cells. Ann Biomed Eng, 34, 879–92.

Hubert, M. & Van Der Veeken, S. 2008. Outlier detection for skewed data. Journal of Chemometrics, 22, 235–246.

Inaishi, J., Saisho, Y., Sato, S., Kou, K., Murakami, R., Watanabe, Y., Kitago, M., Kitagawa, Y., Yamada, T. & Itoh, H. 2016. Effects of obesity and diabetes on α- and β-cell mass in surgically resected human pancreas. J Clin Endocrinol Metab, 101, 2874–82.

Jones, C. N., Pei, D., Staris, P., Polonsky, K. S., Chen, Y. D. & Reaven, G. M. 1997. Alterations in the glucose-stimulated insulin secretory dose-response curve and in insulin clearance in nondiabetic insulin-resistant individuals. J Clin Endocrinol Metab, 82, 1834–8.

Kubota, H., Noguchi, R., Toyoshima, Y., Ozaki, Y., Uda, S., Watanabe, K., Ogawa, W. & Kuroda, S. 2012. Temporal coding of insulin action through multiplexing of the AKT pathway. Mol Cell, 46, 820–32.

Kubota, N., Kubota, T., Itoh, S., Kumagai, H., Kozono, H., Takamoto, I., Mineyama, T., Ogata, H., Tokuyama, K., Ohsugi, M., Sasako, T., Moroi, M., Sugi, K., Kakuta, S., Iwakura, Y., Noda, T., Ohnishi, S., Nagai, R., Tobe, K., Terauchi, Y., Ueki, K. & Kadowaki, T. 2008. Dynamic functional relay between insulin receptor substrate 1 and 2 in hepatic insulin signaling during fasting and feeding. Cell Metab, 8, 49–64.

Lagarias, J., Reeds, J., Wright, M. & Wright, P. 1998. Convergence properties of the Nelder-Mead simplex method in low dimensions. Siam J Optimiz, 9, 112–47.

Lee, C. C., Haffner, S. M., Wagenknecht, L. E., Lorenzo, C., Norris, J. M., Bergman, R. N., Stefanovski, D., Anderson, A. M., Rotter, J. I., Goodarzi, M. O. & Hanley, A. J. 2013. Insulin clearance and the incidence of type 2 diabetes in Hispanics and African Americans: the IRAS Family Study. Diabetes Care, 36, 901–7.

Licko, V. & Silvers, A. 1975. Open-loop glucose-insulin control with threshold secretory mechanism: analysis of intravenous glucose tolerance tests in man. Mathematical Biosciences, 27, 319–332.

Lindsay, J. R., Mckillop, A. M., Mooney, M. H., Flatt, P. R., Bell, P. M. & O’Harte, F. P. 2003. Meal-induced 24-hour profile of circulating glycated insulin in type 2 diabetic subjects measured by a novel radioimmunoassay. Metabolism, 52, 631–5.

Mari, A., Tura, A., Gastaldelli, A. & Ferrannini, E. 2002. Assessing insulin secretion by modeling in multiple-meal tests: role of potentiation. Diabetes, 51 Suppl 1, S221–6.

Marini, M. A., Frontoni, S., Succurro, E., Arturi, F., Fiorentino, T. V., Sciacqua, A., Perticone, F. & Sesti, G. 2014. Differences in insulin clearance between metabolically healthy and unhealthy obese subjects. Acta Diabetol, 51, 257–61.

Matveyenko, A. V., Liuwantara, D., Gurlo, T., Kirakossian, D., Dalla Man, C., Cobelli, C., White, M. F., Copps, K. D., Volpi, E., Fujita, S. & Butler, P. C. 2012. Pulsatile portal vein insulin delivery enhances hepatic insulin action and signaling. Diabetes, 61, 2269–79.

Mizukami, H., Takahashi, K., Inaba, W., Tsuboi, K., Osonoi, S., Yoshida, T. & Yagihashi, S. 2014. Involvement of oxidative stress-induced DNA damage, endoplasmic reticulum stress, and autophagy deficits in the decline of β-cell mass in Japanese type 2 diabetic patients. Diabetes Care, 37, 1966–74.

Neuhauser, M. & Bretz, F. 2001. Nonparametric all-pairs multiple comparisons. Biometric J, 43, 571–80.

Noguchi, R., Kubota, H., Yugi, K., Toyoshima, Y., Komori, Y., Soga, T. & Kuroda, S. 2013. The selective control of glycolysis, gluconeogenesis and glycogenesis by temporal insulin patterns. Mol Syst Biol, 9, 664.

Ohashi, K., Komada, H., Uda, S., Kubota, H., Iwaki, T., Fukuzawa, H., Komori, Y., Fujii, M., Toyoshima, Y., Sakaguchi, K., Ogawa, W. & Kuroda, S. 2015. Glucose homeostatic law: insulin clearance predicts the progression of glucose intolerance in humans. PLoS One, 10, e0143880.

Okuno, Y., Komada, H., Sakaguchi, K., Nakamura, T., Hashimoto, N., Hirota, Y., Ogawa, W. & Seino, S. 2013. Postprandial serum C-peptide to plasma glucose concentration ratio correlates with oral glucose tolerance test- and glucose clamp-based disposition indexes. Metabolism, 62, 1470–6.

Perley, M. J. & Kipnis, D. M. 1967. Plasma insulin responses to oral and intravenous glucose: studies in normal and diabetic sujbjects. J Clin Invest, 46, 1954–62.

Pfeifer, M. A., Halter, J. B. & Porte, D. 1981. Insulin secretion in diabetes mellitus. Am J Med, 70, 579–88.

Picchini, U., De Gaetano, A., Panunzi, S., Ditlevsen, S. & Mingrone, G. 2005. A mathematical model of the euglycemic hyperinsulinemic clamp. Theor Biol Med Model, 2, 44.

Polidori, D. C., Bergman, R. N., Chung, S. T. & Sumner, A. E. 2016. Hepatic and extrahepatic insulin clearance are differentially regulated: results from a novel model-based analysis of intravenous glucose tolerance data. Diabetes, 65, 1556–64.

Polonsky, K. S., Given, B. D., Hirsch, L., Shapiro, E. T., Tillil, H., Beebe, C., Galloway, J. A., Frank, B. H., Karrison, T. & Van Cauter, E. 1988a. Quantitative study of insulin secretion and clearance in normal and obese subjects. J Clin Invest, 81, 435–41.

Polonsky, K. S., Given, B. D. & Van Cauter, E. 1988b. Twenty-four-hour profiles and pulsatile patterns of insulin secretion in normal and obese subjects. J Clin Invest, 81, 442–8.

Rabkin, R. & Kitaji, J. 1983. Renal metabolism of peptide hormones. Miner Electrolyte Metab, 9, 212–26.

Rudovich, N. N., Rochlitz, H. J. & Pfeiffer, A. F. 2004. Reduced hepatic insulin extraction in response to gastric inhibitory polypeptide compensates for reduced insulin secretion in normal-weight and normal glucose tolerant first-degree relatives of type 2 diabetic patients. Diabetes, 53, 2359–65.

Saisho, Y., Kou, K., Tanaka, K., Abe, T., Kurosawa, H., Shimada, A., Meguro, S., Kawai, T. & Itoh, H. 2011. Postprandial serum C-peptide to plasma glucose ratio as a predictor of subsequent insulin treatment in patients with type 2 diabetes. Endocr J, 58, 315–22.

Sano, T., Kawata, K., Ohno, S., Yugi, K., Kakuda, H., Kubota, H., Uda, S., Fujii, M., Kunida, K., Hoshino, D., Hatano, A., Ito, Y., Sato, M., Suzuki, Y. & Kuroda, S. 2016. Selective control of up-regulated and down-regulated genes by temporal patterns and doses of insulin. Sci Signal, 9, ra112.

Sato, H., Terasaki, T., Mizuguchi, H., Okumura, K. & Tsuji, A. 1991. Receptor-recycling model of clearance and distribution of insulin in the perfused mouse liver. Diabetologia, 34, 613–21.

Sedaghat, A. R., Sherman, A. & Quon, M. J. 2002. A mathematical model of metabolic insulin signaling pathways. Am J Physiol Endocrinol Metab, 283, E1084–101.

Seino, S., Shibasaki, T. & Minami, K. 2011. Dynamics of insulin secretion and the clinical implications for obesity and diabetes. J Clin Invest, 121, 2118–25.

Stumvoll, M., Mitrakou, A., Pimenta, W., Jenssen, T., Yki-JÄRvinen, H., Van Haeften, T., Renn, W. & Gerich, J. 2000. Use of the oral glucose tolerance test to assess insulin release and insulin sensitivity. Diabetes Care, 23, 295–301.

Tamaki, M., Fujitani, Y., Hara, A., Uchida, T., Tamura, Y., Takeno, K., Kawaguchi, M., Watanabe, T., Ogihara, T., Fukunaka, A., Shimizu, T., Mita, T., Kanazawa, A., Imaizumi, M. O., Abe, T., Kiyonari, H., Hojyo, S., Fukada, T., Kawauchi, T., Nagamatsu, S., Hirano, T., Kawamori, R. & Watada, H. 2013. The diabetes-susceptible gene SLC30A8/ZnT8 regulates hepatic insulin clearance. J Clin Invest, 123, 4513–24.

Toffolo, G., Breda, E., Cavaghan, M. K., Ehrmann, D. A., Polonsky, K. S. & Cobelli, C. 2001. Quantitative indexes of beta-cell function during graded up and down glucose infusion from C-peptide minimal models. Am J Physiol Endocrinol Metab, 280, E2–10.

Toffolo, G., Campioni, M., Basu, R., Rizza, R. A. & Cobelli, C. 2006. A minimal model of insulin secretion and kinetics to assess hepatic insulin extraction. Am J Physiol Endocrinol Metab, 290, E169–76.

VØLund, A., Polonsky, K. S. & Bergman, R. N. 1987. Calculated pattern of intraportal insulin appearance without independent assessment of C-peptide kinetics. Diabetes, 36, 1195–202.

VØLund, A., Brange, J., Drejer, K., Jensen, I., Markussen, J., Ribel, U., SØRensen, A. R. & Schlichtkrull, J. 1991. In vitro and in vivo potency of insulin analogues designed for clinical use. Diabet Med, 8, 839–47.

